# The cell polarity proteins Boi1p and Boi2p stimulate vesicle fusion at the plasma membrane of yeast cells

**DOI:** 10.1101/144360

**Authors:** Jochen Kustermann, Yehui Wu, Lucia Rieger, Dirk Dedden, Tamara Phan, Paul Walther, Alexander Dünkler, Nils Johnsson

**Author notes:** Contact: Nils Johnsson, Institute of Molecular Genetics and Cell Biology, Department of Biology, Ulm University, James-Franck-Ring N27, D-89081 Ulm, Germany. equal contribution (Jochen Kustermann, Yehui Wu).

## Abstract

Eukaryotic cells can direct secretion to defined regions of their plasma membrane. These regions are distinguished by an elaborate architecture of proteins and lipids that are specialized to capture and fuse post-Golgi vesicles. Here we show that the proteins Boi1p and Boi2p are important elements of this area of active exocytosis at the tip of growing yeast cells. Cells lacking Boi1p and Boi2p accumulate secretory vesicles in their bud. The essential PH domains of Boi1p and Boi2p interact with Sec1p, a protein required for SNARE complex formation and vesicle fusion. Sec1p loses its tip localization in cells depleted of Boi1p and Boi2p but can partially compensate for their loss upon overexpression. The capacity to simultaneously bind phospholipids, Sec1p, multiple subunits of the exocyst, Cdc42p, and the module for generating active Cdc42p identify Boi1p and Boi2p as essential mediators between exocytosis and polar growth.

**Summary statement:** A novel protein complex connects vesicle fusion with Cdc42p activation. Genetic and protein interaction data suggest that its central members Boi1p and Boi2p chaperone the formation of the docking complex.

## Introduction

The fusion of a secretory vesicle with the plasma membrane (PM) can be conceptually divided into the four discrete steps of vesicle trafficking, membrane tethering, docking and the actual fusion of the two membranes. Each step is thought to require the elaborate and coordinated activities of a suite of proteins whose identities, specific roles and mechanisms of actions are still the subject of intense studies (Südhof and Rothman, 2009; Rizo and Südhof, 2012).

In many eukaryotic cells vesicle fusion does not occur evenly over the entire PM but only at certain areas to promote active cell expansion and the preferred release of signaling- and extracellular matrix molecules. In the budding yeast *Saccharomyces cerevisiae* polar growth is achieved by directing post-Golgi vesicles to the tip of the cell. The RhoGTPase Cdc42p, the master regulator of polar growth in yeast and other eukaryotes, stays at the top of the cascade of proteins that initiates and maintains the polarity of the cell (Bi and Park, 2012). Active Cdc42p (Cdc42_GTP_) is concentrated preferentially at the membrane of the growing tip and enables polarized secretion by at least two interdependent mechanisms. Cdc42_GTP_ activates the yeast formin Bni1p to stimulate the outgrowth of actin cables from this site (Evangelista et al., 1997). Post-Golgi vesicles then ride on these cables towards the cell tip (Donovan and Bretscher, 2015; Donovan and Bretscher, 2012). Here Cdc42_GTP_ binds and possibly activates Exo70p and Sec3p, two members of the exocyst (Zhang et al., 2001 a; Yamashita et al., 2010; Baek et al., 2010; Guo et al., 2001; Adamo et al., 2001). The exocyst belongs to the catcher family of protein complexes that tether incoming vesicles to their target membranes (Yu and Hughson, 2010; Heider et al., 2016). Its eight subunits Sec3p, Sec5p, Sec6p, Sec8p, Sec10p, Sec15p, Exo70p, and Exo84p were first discovered in yeast and later found to perform similar conserved functions during exocytosis in other eukaryotes as well (TerBush and Novick, 1995; TerBush et al., 1996; Yu and Hughson, 2010). Sec10p and Sec15p are suggested to make contact with the membrane of the post-Golgi vesicle whereas Sec3p and Exo70p bind active Cdc42p, and phospholipids at the PM (Wiederkehr et al., 2003; Guo et al., 1999; Zhang et al., 2001 b; Yamashita et al., 2010; Wu et al., 2010; Wu et al., 2008; Boyd et al., 2004; Picco et al., 2017; He et al., 2007).

After vesicle tethering is accomplished the soluble SM protein Sec1p stimulates the formation of an inter-membrane docking complex between the membrane-bound v- and t-SNAREs (soluble N-ethylmaleimide-sensitive-factor attachment receptor) Snc1p, Sso1/2p, and Sec9p (Morgera et al., 2012; Hashizume et al., 2009). A rearrangement of the components of this docking complex, and/or the release of inhibitory factors subsequently initiates membrane fusion (Südhof and Rothman, 2009).

In addition to its role in vesicle trafficking and tethering, Cdc42_GTP_ might stimulate the assembly of a protein scaffold that organizes the sequence of the individual steps of exocytosis at the cortex of the bud (Liu and Novick, 2014). Defining the composition, functions, and architecture of this cortical domain is challenging as it probably forms only transiently during the cell cycle and will coexist with other protein assemblies that simultaneously perform the many further activities at this location.

The homologous proteins Boi1p and Boi2p (Boi1/2p) of *Saccharomyces cerevisiae* are localized below the bud tip and were identified as subunits of a polarity complex including Cdc42p, the GEF for Cdc42p, Cdc24p, and the scaffold protein Bem1p (Bender et al., 1996). As Pob1p, the Boi1/2p-homologue of *Schizosaccharomyces pombe*, was functionally linked to secretion, Boi1/2p are potentially important members of a Cdc42_GTP_-induced active area of secretion at the cell tip (Nakano et al., 2011). Here we characterize Boi1/2p as essential scaffold proteins for exocytosis that assemble Cdc42_GTP_, Bem1p, Cdc24p, the exocyst, and Sec1p into one complex at the PM. By activating Cdc42p and binding directly to the exocyst and Sec1p, this novel complex supports a focused association of the vesicles with the PM and might stimulate the formation of the docking complex.

## Results

### Depletion of Boi1/2p disrupts the fusion of secretory vesicles with the PM

Boi1/2p consist of an N-terminal SH3 domain (SH3_Boi1_; SH3_Boi2_) a central SAM domain and a C-terminal PH domain (PH_Boi1_, PH_Boi2;_ Fig 1A). An extended linker region between the SAM and the PH domain contains the PXXP motif (P) that binds to the second SH3 domain of Bem1p (SH3_2_Bem1_) (Bender et al., 1996). Sporulation analysis of a heterozygous *BOI1*/Δ*boi1; BOI2*/Δ*boi2* strain confirms that the presence of either *BOI1* or *BOI2* is necessary for cell survival (Fig.1B). The Δ*boi1*Δ*boi2* strain can be rescued by the expression of fragments of the proteins harboring either PH_Boi1_ or PH_Boi2_ (Fig.1C) (Bender et al., 1996). To investigate the essential cellular functions of Boi1/2p we created a Δ*boi2 P_GAL1_ BOI1* strain where the shift to glucose medium killed the cells by repressing the expression of Boi1p (Fig.1D). The overexpression of Boi1p is not toxic in cells lacking Boi2p (Bender et al., 1996). We examined Δ*boi2 P_GAL1_ BOI1* cells by transmission electron microscopy (TEM) 12 hours after the shift from galactose-to glucose-containing medium. Δ*boi2*-cells depleted of Boi1p show a massive accumulation of cytosolic vesicles in their buds (Figs.1E, F). The vesicles display a characteristic bilayer structure with variable diameters of in average 90-100 nm (Fig. 1E, G). The phenotype resembles the accumulation of post-Golgi vesicles in the late *sec* mutants (Novick et al., 1980). To gain independent evidence of the nature of these vesicles we repeated the shut-off experiment with a Δ*boi2 P_GAL1_BOI1* strain that additionally expressed a mCherry fusion to Sec4p, the yeast Rab GTPase and marker for post-Golgi vesicles (Guo et al., 1999). Cells depleted of Boi1/2p show a more intense mCherry-Sec4p staining at the bud tip than cells expressing a single copy of Boi1p (Fig 1I). In contrast to wild type cells the mCherry-signals in Δ*boi2 P_GAL1_BOI1*-cells are not confined to the cell tip but trail toward the mother cell (Fig. 1H). Occasionally we observed a bending of the bud. mCherry-Sec4p was concentrated below the new tip (Fig. 1H). We conclude that the miss-localization of mCherry-Sec4p mirrors the accumulation of vesicles in the TEM pictures of Δ*boi2 P_GAL1_BOI1*-cells.

**Figure 1:**
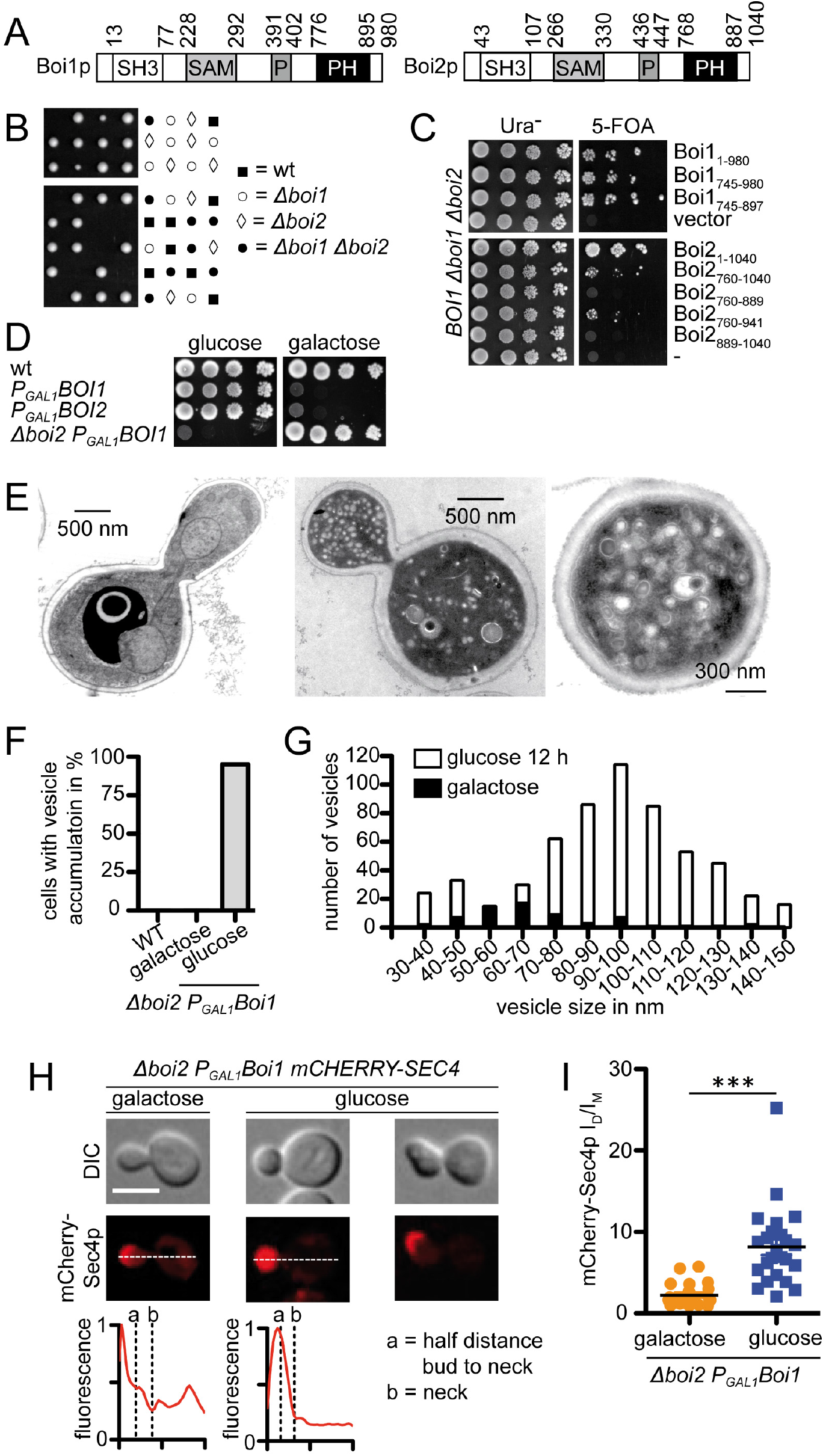
Boi1p and Boi2p are essential for exocytosis. A) Cartoons of the domain structures of Boi1p and Boi2p. SH3= Src homology domain 3. SAM= Sterile alpha motif. P= Bem1p binding motif. PH = Pleckstrin homology domain. B) Sporulation analysis of the diploid *BOI1/Δboi1; BOI2/Δboi2*. Tetrads of the diploid strain were dissected and single spores placed onto YPD medium. C) Δ*boi1* Δ*boi2*-cells expressing *BOI1* from an extra-chromosomal *URA3* plasmid and the indicated Boi1p and Boi2p fragments on a separate centromeric plasmid were spotted in 10-fold serial dilutions onto SD media lacking uracil (left panel), or containing uracil and 5-FOA to kick out the *BOI1-* harbouring URA3-plasmid. D) Wild type-, *P_GAL1_BOI1-, P_GAL1_BOI2-, and Δboi2 P_GAL1_BOI1*-cells were spotted in 10-fold serial dilutions onto media containing galactose (right panel) to fully express or on glucose (left panel) to repress the expression of the *P_GAL1_*-driven genes. E) Δ*boi2 P_GAL1_BOI1*-cells were incubated in galactose (left panel) and shifted to glucose containing media for 12 h (middle and right panel) before processing for TEM. F) Percentage of galactose- or glucose-grown Δ*boi2 P_GAL1_BOI1*-cells that show an accumulation of vesicles in the TEM pictures of their buds. 39< n< 101 cells were analyzed. G) Size distribution of the accumulated vesicles as measured from the TEM pictures in E) and F). H) Fluorescence intensity profiles (lower panel) across the bud of Δ*boi2 P_GAL1_BOI1*-cells expressing mCherry-Sec4p that were either grown in galactose (left panel) or glucose (middle and right panel). Notice the shift in the peak of fluorescence intensity towards the center of the bud in glucose-grown cells. Right panel shows a typical example of a cell with a bent bud. I) Plot of the calculated ratios of the intensities of mCherry-Sec4p in mother and emerging daughters of Δ*boi2 P_GAL1_BOI1*-cells grown in either galactose or glucose. n=23 cells were analyzed. *** = p<0.005. Scale bar 5 μm.

### The PH-domains of Boi1/2p interact with Sec1p

PH_Boi1_ and PH_Boi2_ bind to lipids and active Cdc42p (Bender et al., 1996; Hallett et al., 2002). Both activities are not specifically linked to the accumulation of post-Golgi vesicles. We thus assumed that PH_Boi1/2_ might contact additional ligands that are more directly connected to exocytosis. To identify this hypothetical ligand we searched for new interaction partners of Boi1/2p by a systematic Split-Ubiquitin (Split-Ub) interaction screen using an array of 389 yeast cells, each expressing the N-terminal half of Ubiquitin (N_ub_) as a fusion to a different yeast protein (Müller and Johnsson, 2008; Johnsson and Varshavsky, 1994; Wittke et al., 1999; Hruby et al., 2011). The N_ub_ fusions containing cells were mated with a yeast strain carrying the protein under study as an N-terminal fusion to the C-terminal half of Ubiquitin (C_ub_). C_ub_ is C-terminally extended by an N-degron (R) followed by the uracil-synthesizing enzyme Ura3p (CRU). A diploid strain carrying a pair of interacting N_ub_ and C_ub_ fusions will reconstitute a native-like Ub. The cleaved RUra3 is subsequently degraded and the cells will survive on medium containing 5-Fluoro-orotic acid (5-FOA). 5-FOA kills all other cells with remaining Ura3p activity. The array is enriched in N_ub_ fusions to genes known to be involved in polar growth, secretion, the cytoskeleton, and stress responses (Hruby et al., 2011; Dünkler et al., 2012). As the relevant hits should play essential roles in secretion and should equally interact with both Boi1p and Boi2p in a PH-domain dependent manner, we compared the interaction profiles of the CRU fusions of full length Boi1p and Boi2p with the profiles of their fragments lacking the PH-domains (Boi1_1-733_CRU, and Boi2_1-760_CRU) (Table 1, Fig. S1). N_ub_-Sec1p turned out to be the only fusion to an essential component of the exocytosis pathway whose affinity to Boi1CRU and Boi2CRU substantially decreases upon deletion of their PH domains (Fig. 2A; Table 1; Fig. S1). The Split-Ub analysis of the CRU fusion to the isolated PH domain of Boi1p (PH_Boi1_CRU) suggests a direct binding between PH_Boi1_ and Sec1p or Cdc42p, whereas the newly found interaction between Boi1p and Bud6p is restricted to regions N-terminal to PH_Boi1_ (Fig. 2B, Table 1) (Wittke et al., 1999). We next reconstituted the PH_Boi1/2_/Sec1p complexes with bacterially expressed fusion proteins. Enriched 6xHIS-Sec1p is specifically precipitated by GST-PH_Boi1_ and GST-PH_Boi2_ (Fig. 2C). We used SPLIFF to determine the dynamics of the Boi1p-Sec1p interaction in living cells (Moreno et al., 2013). SPLIFF is a recent modification of the Split-Ub technique. Here the Cub is sandwiched between the auto-fluorescent mCherry and GFP (CCG) (Moreno et al., 2013). Upon the interaction-induced reassociation with a N_ub_ fusion the GFP is cleaved off and rapidly degraded. The subsequent local increase in the ratio of red to green fluorescence indicates where and when the interaction between both proteins took place. Yeast cells expressing Boi1Cherry-C_ub_-GFP (Boi1CCG) were mated under the microscope with cells expressing N_ub_-Sec1p. The N_ub_-induced conversion of Boi1CCG to Boi1CC was subsequently recorded by fluorescence microscopy during the first cell cycle of the generated diploid cells (Fig. 2D) (Moreno et al., 2013; Dünkler et al., 2015). The interaction occurs immediately after cell fusion, later at the incipient bud site and during early bud growth at the tip of the cell. The fast conversion of Boi1CCG is considerably slowed down during the growth of larger buds and during mitosis and cytokinesis (Fig. 2D). The measured interactions are specific, as a N_ub_ fusion to Ptc1p, a protein that does not bind to Boi1p, hardly converts Boi1CCG above background (Fig. 2D).

**Figure 2:**
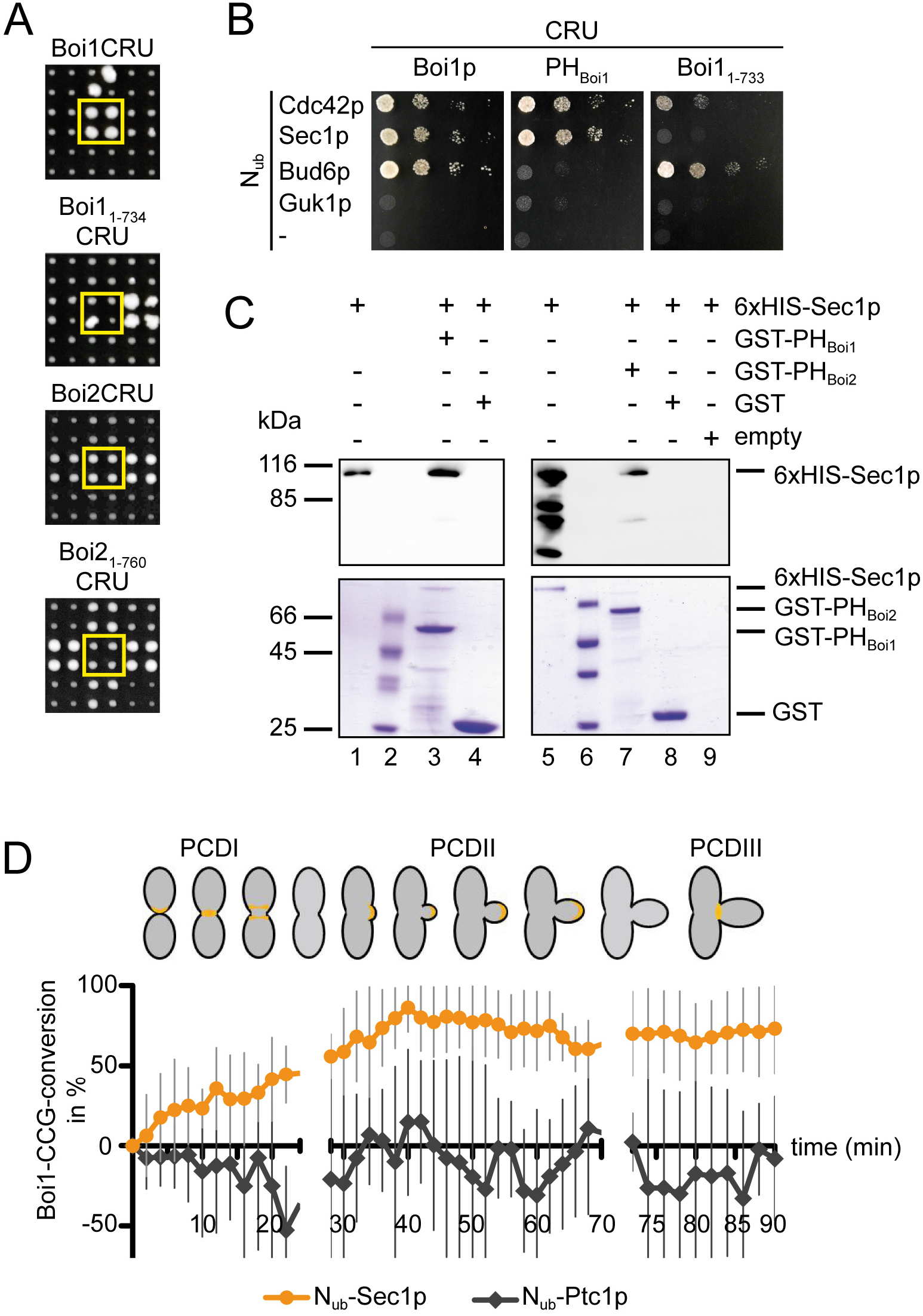
The PH domains of Boi1/2p interact with Sec1p. A) Cut-outs from Split-Ub interaction assays of 389 yeast strains each containing a different N_ub_-fusion and co-expressing from top to bottom: Boi1CRU, Boi1_1-733_CRU, Boi2CRU, or Boi2_1-760_CRU. Shown are quadruplets of each strain grown on medium containing 5-FOA and 50μM Cu for three days (upper two panels), or on 5-FOA and 100μM Cu for two days (lower two panels). Yellow boxes contain the strains co-expressing N_ub_-Sec1p. B) Yeast cells coexpressing the indicated N_ub_ and C_ub_ fusion proteins were spotted in 10 fold serial dilutions on SD medium containing 5-FOA. C) 6xHIS-Sec1p (lanes 1, 5) was incubated with Glutathione-coupled beads pre-treated with GST (lanes 4, 8), with GST fusions to PH_Boi1_ (lane 3), or PH_Boi2_ (lane 7) or with no protein (lane 9). Glutathione-eluates were separated by SDS-PAGE and stained with Coomassie (lower panel) or with anti-HIS antibody (upper panel). Lanes 2, 6: molecular weight marker. D) (A) SPLIFF analysis of Boi1CCG. a-cells expressing Boi1CCG were mated with alpha-cells expressing N_ub_-Sec1p (yellow line), or N_ub_-Ptc1p (grey line). The calculated fractions of converted Boi1CC were plotted against time. Each value represents averages of n=13 independent matings each (error bars, s.e.). Colored regions in the yeast cell cartoons indicate the localization of BOI1CCG and the area used for signal quantification at the respective cell cycle stage (PCD I-III).

**Table 1.** Shared PH domain-dependent interaction partners of Boi1p and Boi2p. The interaction partners of the above CRU constructs were identified by large scale Split-Ub analyses and the presence (+) or absence (−) of interactions recorded in reference to the Boi1p binding partners (Fig. S1)

| CRU<br>N<sub>ub</sub> | Boi1p | Boi1<sub>1-733</sub> | Boi2p | Boi2<sub>1-760</sub> |
| --- | --- | --- | --- | --- |
| Bud6p | + | + | + | - |
| Fks1p | + | + | + | - |
| Boi1p | + | + | + | - |
| Hof1p | + | - | - | - |
| Tpk1p | + | + | + | + |
| Cbk1p | + | + | + | + |
| Epo1p | + | + | + | - |
| Boi2p | + | + | + | + |
| Cdc24 | + | + | + | + |
| Bem1p | + | + | + | + |
| Rga2p | + | + | + | - |
| Rsr1p | + | - | + | - |
| Sec4p | + | + | + | - |
| Nba1p | + | + | + | + |
| Ydl173Wp | + | + | + | + |
| Ras1p | + | - | + | - |
| Sec1p | + | - | + | - |
| Rho4p | + | - | + | - |
| Cdc42p | + | + | + | - |

### Overexpression of Sec1p partially compensates for the depletion of Boi1/2p

Our analysis suggests that PH_Boi1/2_ might stimulate vesicle fusion by recruiting Sec1p to a phospholipid- and Cdc42_GTP_-enriched patch of the PM. If correct, increasing concentrations of Sec1p might compensate for the depletion of Boi1/2p. We inserted the methionine-repressible *P_MET17_* promoter in front of the genomic loci of *SEC1*, of selected members of the exocyst and of Sec4p. The *P_MET17_* promoter is repressed at high methionine concentrations, moderately active at 70 μM and fully active in medium lacking methionine. Of all tested proteins only the overexpression of Sec1p rescues the growth of the Δ*boi2 P_GAL1_BOI1*-cells on glucose medium (Fig. 3A). Similarly, a plasmid-borne GFP fusion to *SEC1* under control of the *P_MET17_* promoter enables growth of Δ*boi2 P_GAL1_BOI1*-cells in medium containing glucose but no methionine (Fig. 3B). TEM analysis of these cells shows a clear reduction in the amount of accumulated vesicles when compared to the same cells but grown in the presence of methionine, or when compared to Δ*boi2 P_GAL1_BOI1*-cells expressing a GFP containing plasmid without *SEC1* (Fig. 3C). Quantifying the fluorescence intensity of mCherry-Sec4p indirectly confirms that overexpression of GFP-Sec1p partially suppresses vesicle accumulation upon depletion of Boi1/2p (Fig. 3D). Overexpression of Sec1p does however not reduce the occurrence of bent buds that are reproducibly encountered in Δ*boi2 P_GAL1_BOI1*-cells (Fig. 1H). The cellular distribution of accumulated vesicles remains less centered to the tip (Fig. 3E). This effect is not encountered in other secretion-defective mutants as for example *sec1-1* cells never display bent buds at either restrictive or non-restrictive temperature (Fig. 3E) (Novick and Schekman, 1979). Sec1p associates with the SNARE complex and supports its assembly and function during vesicle fusion (Morgera et al., 2012; Hashizume et al., 2009). Boi1/2p might stimulate the formation of the SNARE complex by presenting Sec1p to one of its subunits or to a certain intermediate that accumulates during the assembly process. The hypothesis predicts that similar to Sec1p an increase in cellular concentration of this subunit/intermediate alleviates the loss of Boi1/2p. Indeed overexpression of the T-SNARE Sso1p but not of the V-SNARE Snc1p supports growth of Δ*boi2 P_GAL1_BOI1*-cells on glucose (Fig. 6F).

**Figure 3:**
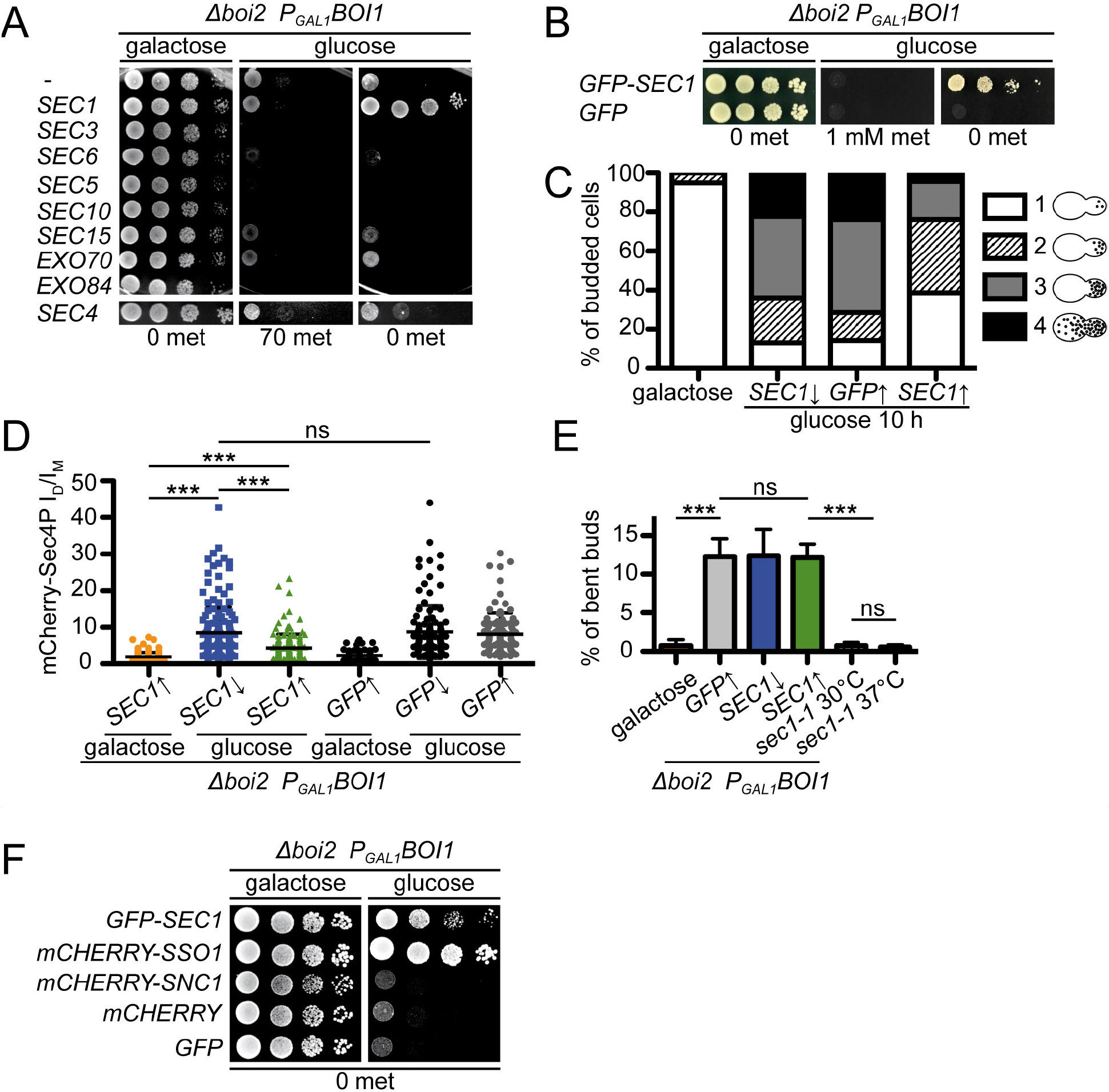
Overexpression of Sec1p rescues the growth of Boi1/2p-depleted cells. A) Δ*boi2 P_GAL1_BOI1*-cells expressing the indicated genes from the integrated *P_MET17_-promoter* were spotted in serial dilutions onto media moderately expressing the indicated genes and expressing (left panel) or repressing (middle panel) Boi1p, and onto media repressing the expression of Boi1p and inducing the expression of the indicated genes (right panel). B) Δ*boi2 P_GAL1_BOI1*-cells expressing *P_Met17_GFP-SEC1* or *P_Met17_GFP* were spotted onto media inducing the expression of Boi1p (left panel) and onto media repressing the expression of Boi1p and either repressing (middle panel) or overexpressing (right panel) GFP-Sec1p or GFP. C) TEM analysis of cells from above. Vesicles were counted in cells grown in galactose, or after a 10 hour-incubation in glucose, and either overexpressing or repressing GFP-Sec1p or GFP. D) Quantification of the fluorescence profiles of mCherry-Sec4p in Δ*boi2 P_GAL1_BOI1*-cells co-expressing *P_Met17_GFP-SEC1* or *P_Met17_GFP*, and in sec1-1-cells at 30°C, or at 37°C. Expression of Boi1p and *P_Met17_GFP-SEC1* or *P_Met17_GFP* was adjusted as in B). The analysis of the mCherry fluorescence was performed as in 1H). A minimum of n > 29 cells in three independent experiments were analyzed for each condition. E) Quantification of bent buds. Cells were grown under identical conditions as in C). n > 28 cells were counted in each of three experiments. ns= not significant;*** = p<0.0005. F) As in B) but additionally expressing the mCherry fusions of Sso1p and Snc1p.

### Binding to Cdc42p is not essential for Boi1/2p’s role in vesicle fusion

Mutations in PH_Boi1/2_ that are known to interfere with phospholipid binding disrupt the essential functions of the proteins (see also Fig. 4E) (Hallett et al., 2002). To find out whether binding to Cdc42 is equally important we first thought to identify residues in PH_Boi1_ that are required for complex formation with active Cdc42p. Homology modeling of the complex between PH_Boi1_ and Cdc42p using the solved structure of human Exo84 bound to human Rall_GTP_ as template identified residues at position R827, L829, and T894 on PH_Boi1/2_ as potentially important for binding active Cdc42p (https://honiglab.c2b2.columbia.edu/PrePPI/html/tmp/P38041_P19073/model.htm;) (Fig. 4A) (Jin et al., 2005). We replaced all three residues with alanines and could demonstrate that a GST fusion of the resulting triple mutant PH_Boi1RLT_ no longer measurably interacts with the constitutively active Cdc42_G12V_ (Fig. 4B). However, GST-PH_Boi1RLT_ still co-sediments with phospholipid vesicles (Fig. 4C), and precipitates purified 6xHIS-Sec1p with comparable efficiency as the GST-fusions to wild type PH_Boi1_, to PH_Boi2_, or to a mutant of Boi1p that lost the ability to interact with lipids (PH_Boi1KKTK_) (Fig. 4D) (Hallett et al., 2002). GFP-Boi1_RLT_ but not GFP-Boi1_KKTK_ supports growth of Δ*boi2 P_GAL1_BOI1*-cells on glucose (Fig. 4E). As both proteins are expressed at comparable levels, we conclude that binding Cdc42p in contrast to binding phospholipids is not essential for Boi1/2p’s role in exocytosis (Fig. S2).

**Figure 4:**
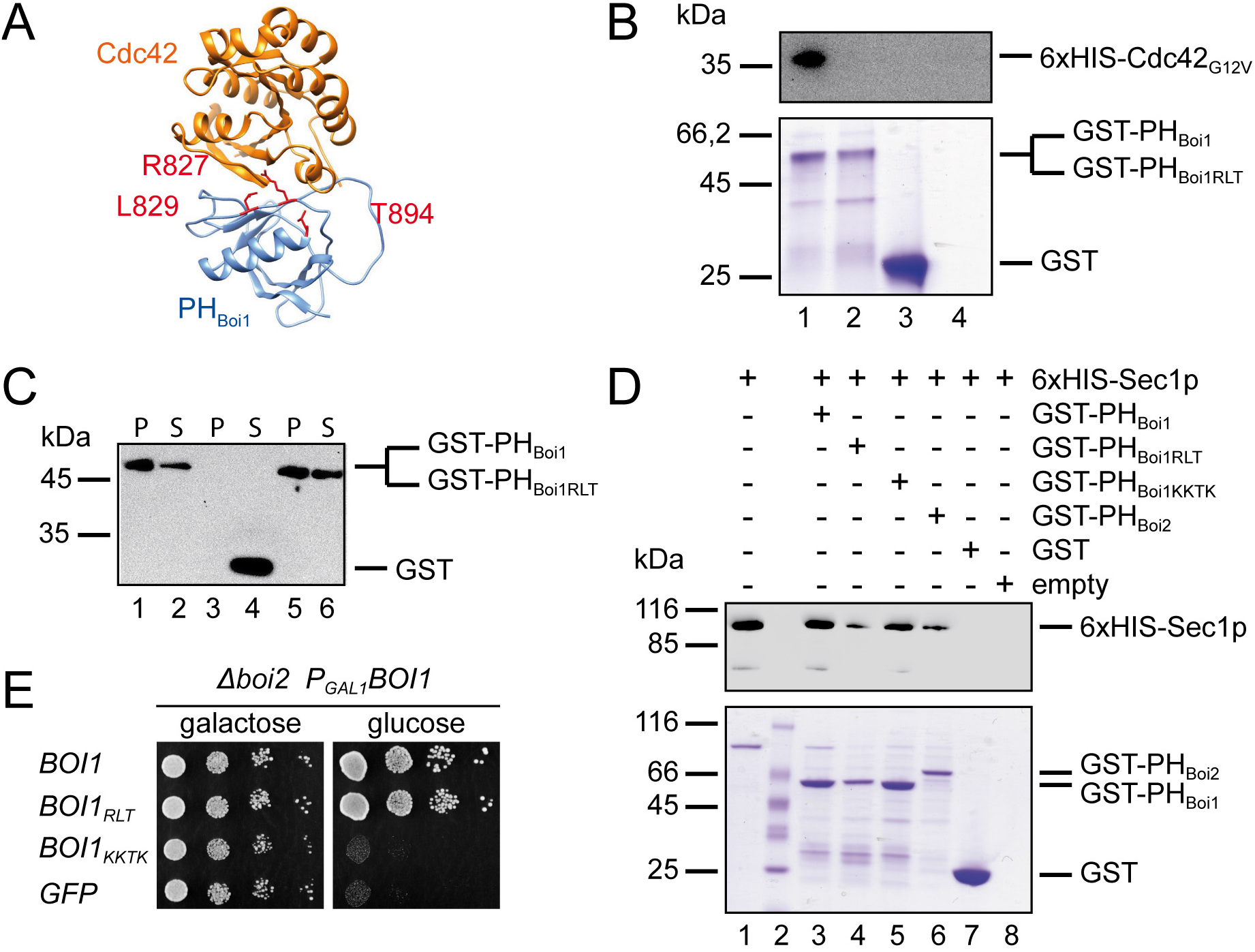
Binding to lipid but not to Cdc42p is essential for the functions of Boi1/2p. A) Proposed structure of the PH_Boi1/2_/Cdc42p complex. Highlighted in red are the altered residues in PH_Boi1RLT_. B) Enriched 6xHis-Cdc42_G12V_ was incubated with Glutathione-coupled beads that were pre-treated with either GST-PH_Boi1_ (lane 1), GST-PH_Boi1RLT_ (lane 2), GST (lane 3), or no protein (lane 4). Glutathione-eluates were separated by SDS-PAGE and either stained with Coomassie-blue (lower panel) or with anti-HIS antibody (upper panel). C) Liposomes incubated with GST-PH_Boi1_ (lanes 1, 2), GST (lanes 3, 4) or GST-PH_Boi1RLT_ (lanes 5, 6). Supernatants (lanes 2, 4, 6) and pellets (lanes 1, 3, 5) were separated by SDS-PAGE and stained with anti-GST antibody. D) Enriched 6xHis-Sec1p (lane 1) was incubated with either immobilized GST-PH_Boi1_ (lane 3), GST-PH_Boi1RLT_ (lane 4), GST-PH_Boi1KKTK_ (lane 5), GST-PH_Boi2_ (lane 6), GST (lane 7), or empty beads (lane 8). Glutathione-eluates were separated by SDS-Page and stained with Coomassie (lower panel) or anti-HIS antibody (upper panel). Lane 2: Molecular weight marker. E) Δ*boi2 P_GAL1_BOI1*-cells expressing GFP, or GFP fusions to Boi1p, Boi1_RLT_, or Boi1_KKTK_ were spotted in 10-fold serial dilutions onto SD medium containing either galactose or glucose.

### Binding to lipid is not sufficient for Boi1/2p’s role in vesicle fusion

Sec1p overexpression suppresses the effects of Boi1/2p depletion (Fig. 3). This observation strongly suggests that the interaction between Sec1p and PH_Boi1/2_ is functionally relevant. To further test this hypothesis we searched for Boi1/2p homologues that might still bind lipid and Cdc42p but not Sec1p of S. *cerevisiae*. Sequence homologues of Boi1/2p can be found in closely related yeasts, but also in Schizosaccharomyces *pombe* and more distant fungi like *Aspergillus nidulans* (Nakano et al., 2011). Alignment of the PH domains of the Boi-proteins from *S.cerevisiae* (PH_Boi1_), *S.pombe* (PH_SpBoi_), and *A. nidulans* (PH_AnBoi_) identifies the conserved lipid binding motif and the binding motif for Cdc42p in the sequences of both species (Fig. 5A). However, although detected at comparable levels in protein extracts, only Δ*boi2 P_GAL1_BOI1*-cells cells expressing PH_SpBoi_ but not cells expressing PH_AnBoi_ survive on glucose-containing medium (Fig. 5B). Co-precipitation experiments prove that both PH domains show similar binding to phospholipid and Cdc42_G12V_ (Figs. 5C, D, S3A, B). We thus tested whether binding to Sec1p might distinguish the complementing PH_SpBoi_ from the non-complementing PH_AnBoi_. Pull down experiments show a significantly reduced affinity of GST-PH_AnBoi_ for His_6_-Sec1p whereas the complementing PH_SpBoi_ seems to bind 6xHis-Sec1p slightly stronger than PH_Boi1_ (Fig. 5E, see also Fig. S3C, D).

**Figure 5:**
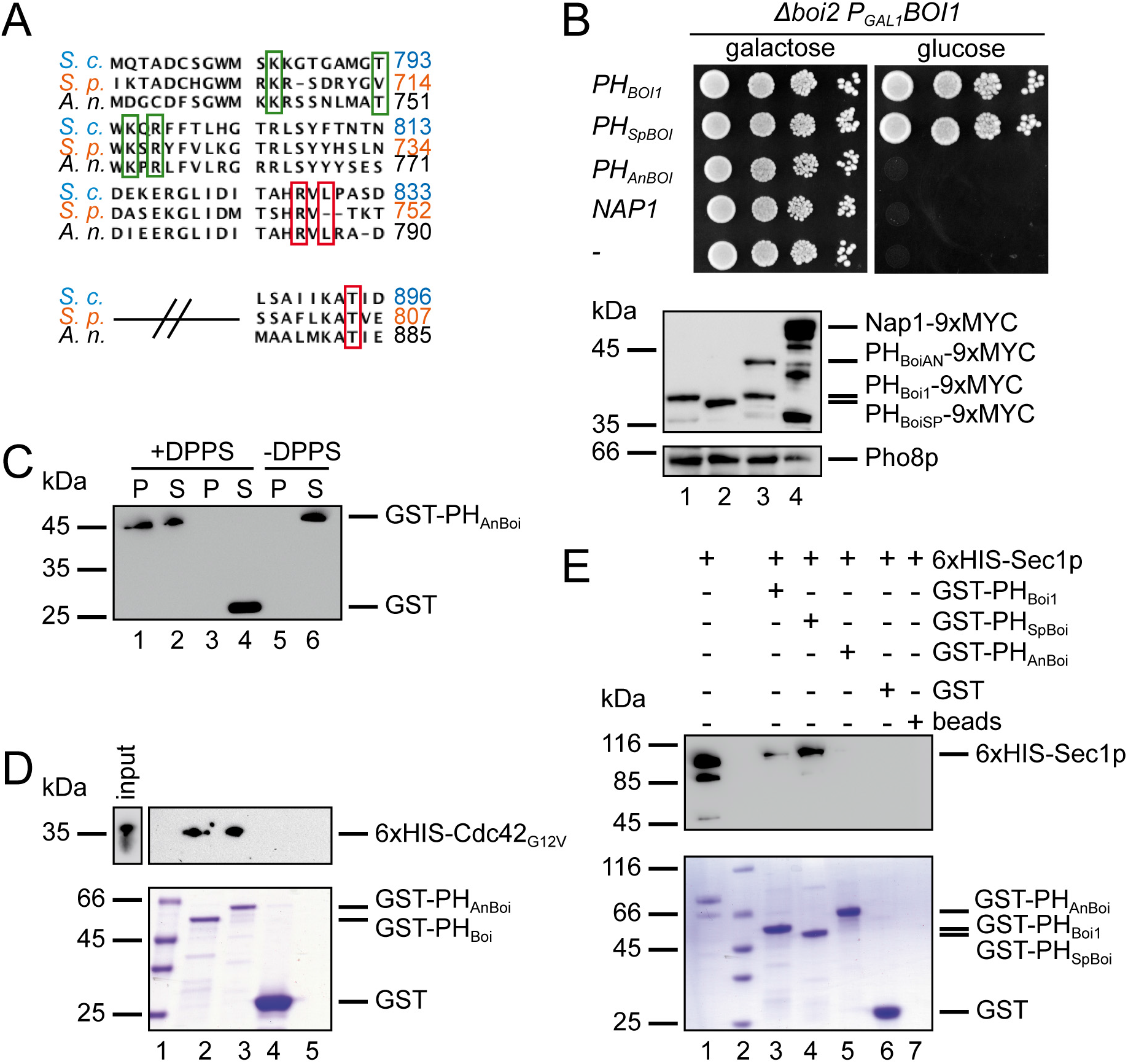
The PH domain of Boi1p from *A. nidulans* does not complement the loss of *BOI1/2* in *S.cerevsiae*. A) Sequence alignment of the PH domains of Boi1p from *S.cerevisiae, S.pombe* and *A. nidulans*. Boxed in green and red are residues critical for binding lipid (green) or Cdc42p (red). B) Upper panel: Δ*boi2 P_GAL1_BOI1-cells* expressing the 9xMYC tagged PH domains of the indicated fungi, the yeast protein Nap1p, or an empty plasmid were spotted in serial dilutions onto media containing galactose to stimulate, or glucose to inhibit the expression of the full-length Boi1p. Lower panel: Protein extracts of Δ*boi2 P_GAL1_BOI1-cells* expressing 9xMYC-PH_Boi1_ (lane 1), 9xMYC-PH_SpBoi_ (lane 2), 9xMYC-PH_AnBoi_ (lane 3), 9xMYC-Nap1 (lane 4) or no MYC-labeled protein (lane 5) were separated by SDS-PAGE and stained with anti-MYC antibody. The extracts were additionally probed with anti-Pho8p antibodies (lower panel). C) Liposomes (DPPS) (lanes 1-4) or buffer without liposomes (lanes 5, 6) were incubated with GST-PH_AnBoi_ (lanes 1, 2, 5, 6) or GST (lanes 3, 4) and spun down. Fractions of pellet (P) and supernatant (S) were separated by SDS-PAGE and stained with anti-GST antibody. D) Enriched 6xHis-Cdc42_G12V_ was incubated with Glutathione-coupled beads either pretreated with GST-PH_Boi1_ (lane 2), GST-PH_AnBoi_ (lane 3), GST (lane 4), or no protein (lane 5). Glutathione-eluates were stained with anti-HIS antibodies after SDS-PAGE and transfer onto nitrocellulose. Lane 1: Molecular weight marker. E) Enriched 6xHis-Sec1p (lane 1) was incubated with Glutathione-coupled beads either pre-treated with GST-PH_Boi1_ (lane 3), GST-PH_SpBoi_ (lane 4), GST-PH_AnBoi_ (lane 5), GST (lane 6), or no protein (lane 7). Glutathione-eluates were separated by SDS-GAGE and stained with anti-HIS antibodies (upper panel) or with Coomassie (lower panel). Lane 2: Molecular weight marker.

### Boi1/2p assists in focusing Sec1p to the cell tip

The hammerhead morphologies occasionally displayed by Δ*boi2 P_GAL1_BOI1*-cells after switch to glucose suggests that Boi1/2p keep the area of exocytosis focused to the front end of the cells (Figs. 1H, 6A). In agreement, the depletion of Boi1/2p shifts the distribution of GFP-Sec1p from its polar location at the bud tip more towards the sides of the bud, to the tips of bent buds, or occasionally into mother cells (Fig. 6A, B). To find out whether Boi1/2p reciprocally require a functional Sec1p for their correct cortical localization, we shifted *sec1-1ts* cells co-expressing mCherry-Sec4p and Boi1-GFP to their restrictive temperature (Novick and Schekman, 1979). Although the accumulation of mCherry-Sec4p becomes clearly apparent after the temperature shift, Boi1-GFP remains predominantly attached to the cortex (Fig. 6C). Latrunculin A treatment disrupts the actin cytoskeleton of yeast cells and thus the traffic of post-Golgi vesicles to the bud (Ayscough et al., 1997). As the drug does not measurably affect the cortical localization of Boi1- or Boi2-GFP (Fig. 6D, E), we conclude that Boi1/2p do not reach the PM through vesicular traffic but instead are part of the structure that receives and processes vesicles at the cortex.

**Figure 6:**
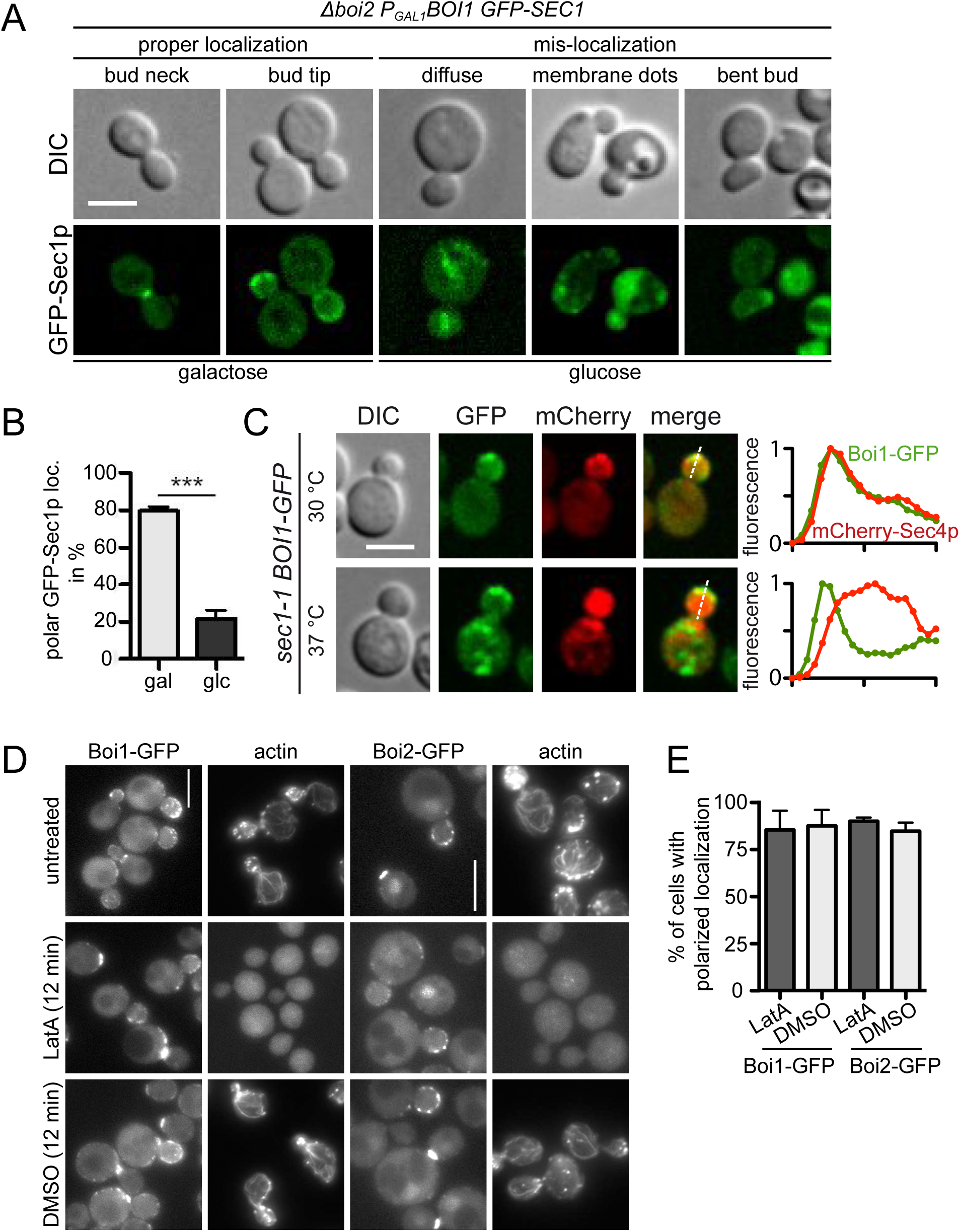
Sec1p requires Boi1/2p for proper localization. A) Fluorescence microscopy of Δ*boi2 P_GAL1_BOI1-cells* expressing GFP-Sec1p either grown in galactose to induce or grown for 10 hours in glucose to repress the expression of Boi1p. B) Quantification of polar GFP-Sec1p localization in cells as in A). A minimum of n > 153 cells were counted in each of three independent experiments. C) Fluorescence microscopy of *sec1-1*-cells co-expressing Boi1-GFP with mCherry-Sec4p at 30°C (upper panels) or after a 3 h incubation at the restrictive temperature of 37°C (lower panels). The fluorescence intensities of the proteins are plotted across the buds in the right upper and lower panels. D) GFP fusions to Boi1p, or Boi2p, or actin-fibers and -patches were visualized either directly or with phalloidin-rhodamine in untreated cells (upper panels), after treatment with latrunculin A (middle panels), or DMSO (lower panels). E) Quantifications of Boi1-GFP, and Boi2-GFP localizations in cells shown in D). A minimum of n= 65 cells were inspected in each of two experiments. Scale bars 5 μm.

### Boi1/2p bind the exocyst

Boi1/2p bind Bem1p and through Bem1p indirectly Cdc24p (Table 1) (McCusker et al., 2007). The interaction sites for Bem1p are located N-terminally to the PH domains of Boi1/2p (Fig. 1A). The observation that the expression of the isolated PH_Boi1/2_ or of Boi1_RTK_ can rescue the loss of Boi1/2p questions the functional significance of the Boi1/2p mediated connection between exocytosis and Cdc42p-generated polarity. To uncover any potential redundancy that might exist within this connection, we created deletions of the N-terminal 131 or 216 residues of *BEM1* (*bem1_Δ1-131_, bem1_Δ1-216_*) in a Δ*boi1*Δ*boi2* strain ectopically expressing PH_Boi1_. These deletions do not cause a severe growth defect in cells expressing the full length Boi2p but are not tolerated in the strain harboring only PH_Boi1_ (Fig. 7A). The effect is specific for the *bem1* alleles as the removal of some of the other newly discovered binding partners of Boi1/2p had no measurable effect on the growth of the PH_Boi1_-only expressing cells (Fig. 7A). Deletion of the first 138 residues removes the N-terminal SH3 domain of Bem1p that binds to the exocyst subunit Sec15p (France et al., 2006). The lack of this domain in otherwise wild type cells only slightly impairs Sec15p targeting and exocyst complex formation (France et al., 2006). Accordingly, the strong effect of *bem1_Δ1-131_* and consequently of *bem1_Δ1-216_* on the growth of *PH_boi1_*-cells reveals a functional redundancy between *BEM1* and *BOI1/BOI2* that might be explained by a direct physical connection between the exocyst and Boi1/2p. We thus screened members of the exocyst except *SEC10* as CRU fusions against our N_ub_-array and discovered interaction signals between N_ub_-Boi2p and all tested subunits of the exocyst. N_ub_-Boi1p interacts with a smaller subset of the exocyst proteins in our Split-Ub analysis (Fig. 7B; Table S1, Fig. S4). Tonikian at al. predicted for SH3_Boi1_ a binding motif in the sequence of Exo84p, and for SH3_Boi1_ and SH3_Boi2_ a common binding motif in the sequence of Sec3p (Tonikian et al., 2009). Indeed, a N_ub_ fusion to a fragment of Boi1p that lacks SH3_Boi1_ (N_ub_-Boi1_70-980_) fails to interact with Exo84CRU whereas the Split-Ub based interaction signal between Boi2p and Exo84p depends on its PH-but not on its SH3 domain (Fig. 7C). N_ub_-Boi1p interacts in an SH3-dependent manner with the CRU fusions to peptides harboring the prospective binding sites of Sec3p or Exo84p (Fig. 7C). In contrast, N_ub_-Boi2p does not interact or only very weakly with the same peptides (Fig. 7C). Pull-down analysis with the bacterially expressed and enriched GST fusions to SH3_Boi1_ or SH3_Boi2_ confirms that SH3_Boi1_ binds directly whereas SH3_Boi2_ does not bind to either of the peptides (Figs. 7D, S5A). The measured Kd of approximately 1μM for the SH3_Boi1_/ 6xHIS Exo84_121-141_-AGT complex is typical for the weak affinities by which SH3 domains bind their targets (Fig. S5D) (Gorelik and Davidson, 2012).

**Figure 7:**
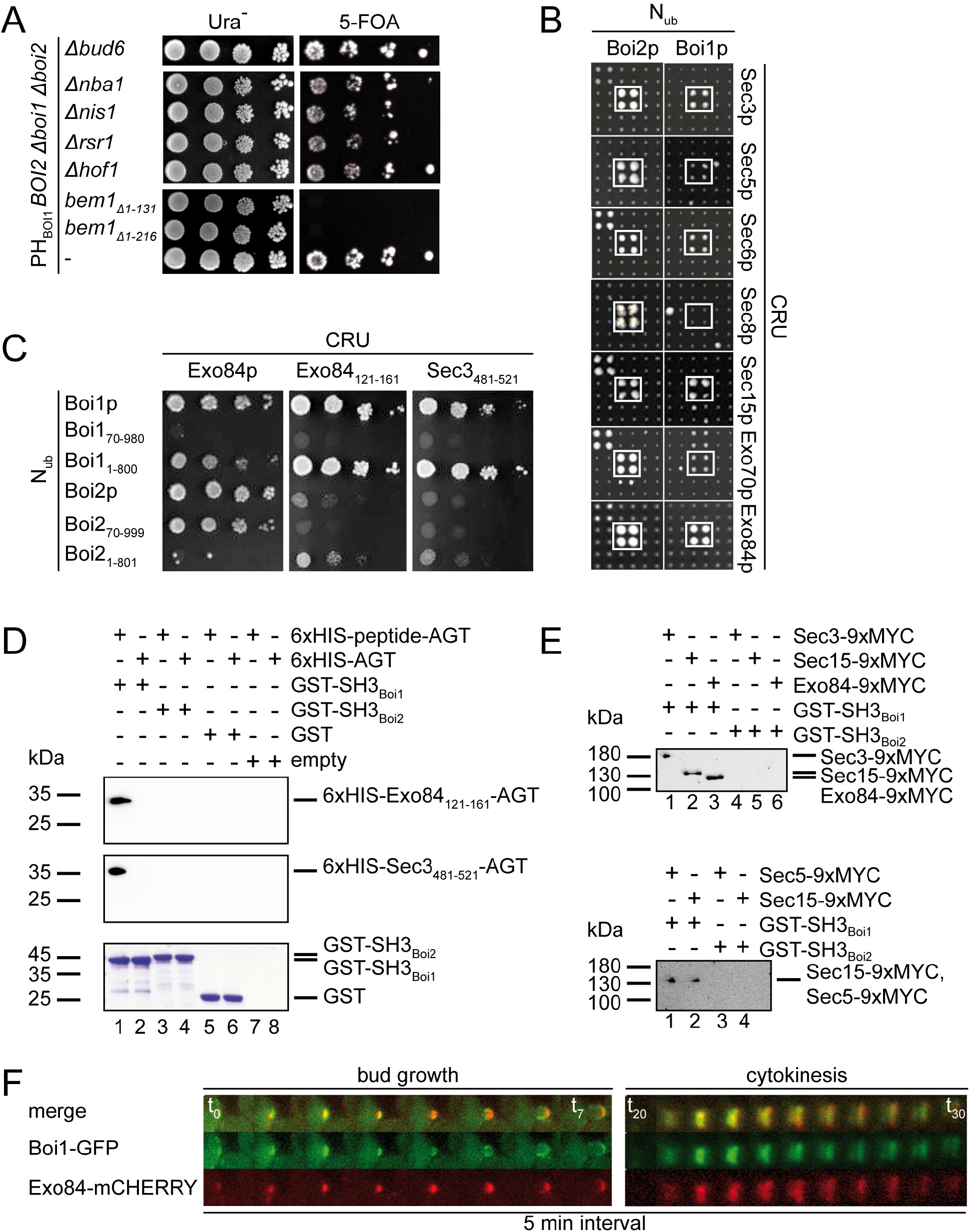
Boi1p and Boi2p interact with the exocyst. A) Δ*boi1*Δ*boi2*-cells carrying the indicated deletions of genes or fragments of genes and expressing *PH_Boi1_* and *BOI2* on different centromeric plasmids were spotted in serial dilutions on SD media containing uracil (left panel), or uracil and 5-FOA (right panel) to kick out the BOI2-containing *URA3* vector. B) Cut-outs from large-scale Split-Ub interaction assays of CRU fusions to members of the exocyst after three days of growth on medium containing 5-FOA and 50μM Cu. White boxes contain the strains co-expressing N_ub_-Boi2p (left panels) or N_ub_-Boi1p (right panels). C) Cells co-expressing the indicated N_ub_ fusions together with CRU fusions to Exo84p, or to peptides derived form Exo84p and Sec3p were spotted in serial dilutions on media containing 5-FOA and 50μm Cu. D) Purified 6xHis-AGT (lanes 2, 4, 6, 8) or 6xHis-AGT fusions to peptides of Exo84p (upper panel: lanes 1, 3, 5, 7) or Sec3p (middle panel: lanes 1, 3, 5, 7) were incubated with immobilized GST (lanes 5, 6) or with immobilized GST fusions to SH3_Boi1_ (lanes 1, 2), SH3_Boi2_ (lanes 3, 4) or with untreated beads (lanes 7, 8). Glutathione-eluates were stained after SDS-PAGE with anti-HIS antibodies or Coomassie. E) Extracts of yeast cells expressing 9xMYC fusions to Sec3p (upper panel: lanes 1, 4) to Sec15p (upper panel: lanes 2, 5; lower panel: lanes 2, 4) to Exo84p (upper panel: lanes 3, 6), or to Sec5p (lower panel: lanes 1, 3) were incubated with immobilized GST-SH3_Boi1_ (upper panel: lanes 1-3; lower panel: lanes 1, 2) or GST-SH3_Boi2_ (upper panel: lanes 4-6; lower panel: lanes 3, 4). Glutathione-eluates were stained as in D). F) Time-lapse analysis of a yeast cell coexpressing Boi1-GFP and Exo84-mCherry. Upper panel shows the merge of the GFP-(middle panel) and the mCherry-channel (lower panel) of pictures from the cell during bud growth (left panel) and from the bud neck during mitosis (right panel).

Using GST-SH3_Boi1_ as bait we enriched from yeast extracts the direct binding partners Sec3p and Exo84p and the exocyst subunits Sec5p, and Sec15p. As the latter do not display consensus sites for SH3_Boi1_, we assume that Boi1/2p bind indirectly to Sec5p and Sec15p. This strongly suggests that Boi1p associates with the intact exocyst complex (Figs. 7E, S5B, C) (Tonikian et al., 2009). The measured interactions between Boi1p and the exocyst are indirectly confirmed by the co-localization of Exo84-mCherry with Boi1-GFP throughout the cell cycle (Fig. 7F).

## Discussion

Our experiments identify Boi1/Boi2p as central scaffold proteins that bind Cdc42_GTP_, its GEF, phospholipid, the exocyst, and Sec1p at the PM below the tip of the cell. The binding sites for the ligands could be mapped to different domains of Boi1/2p or to different interfaces of the same domains. Although not proven a co-existence of the different ligands in one protein complex is thus likely. The known activities of its members and its location at the cell cortex suggests that this novel protein complex plays an important role in reinforcing the maintenance of polarity through the fusion of post-Golgi vesicles at restricted sites of the PM. Indeed, depletion of Boi1/2p leads to the accumulation of post-Golgi vesicles in the buds of the cells.

The PH domains of Boi1/2p carry the essential activities of the proteins and target them to the cell tip (Hallett et al, 2002). Mutations in PH_Boi_ that reduce lipid binding disturb the polar distribution of the proteins and interfere with their functions (Bender et al., 1996; Hallett et al., 2002). Unexpectedly, the expression of the PH domain of *A. nidulans* does not rescue yeast cells lacking Boi1/2p although displaying the conserved lipid binding signature and binding equally well to lipid and Cdc42_GTP_ as its yeast counterparts. However PH_AnBoi_ displays a much weaker affinity to Sec1p suggesting that the newly discovered interaction between PH_Boi1/2_ and Sec1p performs an important step during exocytosis. Overexpression of Sec1p increases the amount of assembled SNARE complexes *in vivo* (Wiederkehr et al., 2004). It was therefore revealing to discover that a raise in Sec1p concentration also partially compensates for the loss of Boi1/2p. This led us to the conclusion that Boi1/2p assist in a critical step during the ordered formation of the SNARE complex. As the t-SNARE Sso1p also suppresses the depletion of Boi1/2p we argue that this critical step involves chaperoning the Sso1p-Sec1p interaction (Fig. S6). Intriguingly, the N-terminal PH domain of Sec3p binds the active conformation of Sso1/2p and promotes its assembly into the Sec9p-Sso1/2p dimer (Yue et al., 2017). Boi1/2p interact directly with Sec3p and all three proteins bind with their PH domains phospholipids and active Cdc42p (Yamashita et al., 2010). Consequently, this small interaction network will align the two PH domain-specific ligands Sso1/2p and Sec1p into very close proximity to promote their assembly into a Sec9p-Sso1/2p-Sec1p trimer (Fig. S6). How and at which step the incorporation of the v-SNARE Snc1/2p occurs is not clear, especially as Snc1/2p is thought to intervene at multiple stages of this process (Carr et al., 1999; Morgera et al., 2012; Hashizume et al., 2009; Carr and Rizo, 2010). Although experimentally supported by others and us the proposed assembly line remains hypothetical unless *in vivo* data on the efficiency and kinetics of SNARE-complex formation become available (Yue et al., 2017).

Vesicle fusion most likely stimulates the hydrolysis of Cdc42_GTP_ to dissociate the exocyst from the membrane. In addition it dilutes membrane-bound polarity factors and Cdc42_GTP_ through the incorporation of new membranes. The observed robustness by which polar growth is maintained is thus difficult to reconcile with a static connection between the exocyst and Cdc42_GTP_. Cdc24p, the GEF for Cdc42p, binds constitutively to Bem1p and through Bem1p to Boi1/2p (Table 1) (McCusker et al., 2007). As part of the exocyst-Boi1/2p-Sec1p complex, Cdc24p is thus placed next to the site of vesicle tethering and fusion (Fig. S6). This coupling could rapidly restore depleted pools of Cdc42_GTP_ and regain the fusion competence of the PM at this site. The design of this CDC42_GTP_-producing tethering complex guides the exocyst, and Sec1p to the area of highest Cdc42p concentration and conversely attracts the source of active Cdc42p to sites where vesicle tethering and fusion occurs. Both effects might reinforce each other to contribute to a stable axis of growth.

We are still at the beginning of characterizing the protein interaction network that underlies the connection between exocytosis and cellular polarity. Our work suggests that Boi1/2p play an instructive part in assembling this network through binding to the exocyst, Bem1p, and Cdc42p (Table 1). Boi1p, Boi2p, and Bem1p share physical and negative genetic interactions with multiple subunits of the exocyst as well as with each other (Figs.1, 7), (France et al., 2006; Liu and Novick, 2014; Costanzo et al., 2010; Costanzo et al., 2016). The non-overlapping binding sites let us propose that Boi1p, Boi2p and Bem1p can be found in one complex with the exocyst (Fig. S6). Accordingly, the growth of a PH_Boi1_-only expressing strain is dependent on the first SH3 domain of Bem1p that binds to Sec15p. The genetic interactions thus complement our protein interaction network and locate the polarity factors Boi1/2p and Bem1p at the center of a cortical sub-domain that organizes polar secretion and cooperates directly in vesicle tethering and docking. The network is held together by a system of SH3 domain interactions that connects the exocyst subunits Sec15p, Exo84p and Sec3p with Boi1p and Bem1p, and Bem1p with Boi1p and Boi2p (Fig. S6). With a Kd of approximately 1μM the single interactions between SH3_Boi1_ and Exo84p and between SH3_2_Bem1_ and Boi1p or Boi2p are weak (Gorelik and Davidson, 2012). However, Boi1p and Boi2p interact with each other and with themselves, SH3_1_Bem1_ interacts with Sec15p, and all three proteins bind to membranes and active Cdc42p. Once combined in a network each single weak interaction might synergistically contribute to a robust connection between exocytosis and the establishment and maintenance of polarity (Fig. S6).

## Materials and methods

### Construction of plasmids and gene fusions

Gene fusions with the coding sequences for the N-terminal 35 residues of ubiquitin (N_ub_), the C-terminal 41 residues of ubiquitin (C_ub_), GFP carrying a S65T exchange, mCherry, a 9xMYC-, or the Cherry-C_ub_-GFP (CCG) module were performed as described (Hruby et al., 2011; Dünkler et al., 2012; Moreno et al., 2013; Neller et al., 2015). Specifically, *BOI1-GFP, BOI1CRU* or *BOI1CCG* were constructed by genomic in-frame insertion of the *GFP, CRU* or *CCG* module behind the genomic *BOI1* ORF. In brief, a PCR fragment of 840 bp of the 3’ end of *BOI1* without stop codon and containing an *EagI* and a *SalI* restriction site at the 3’ and 5’ end respectively, was cloned in front of the *GFP, CRU* or *CCG* module of a pRS303, pRS306, or pRS304 vector (Sikorski and Hieter, 1989). The obtained vectors were linearized using a single *EcoRI* site in the *BOI1* sequence, and transformed into yeast cells. Colony PCR with diagnostic primer combinations was used to verify the successful integration. C-terminal 9xMYC-fusions of exocyst subunits, *EXO84-mCHERRY*, and all C-terminal *CRU* fusions have been generated accordingly. The native genomic promoter sequence was replaced by *P_MET17_, P_MET17_-GFP*, or *P_GAL1_* through recombination with a PCR fragment generated from the pYM-N35, pYM-N37 or pYM-N22 vectors, and primers containing sequences identical to the respective genomic location at their 5’ ends (Janke et al, 2004). To express CRU fusions from centromeric plasmids, the ORFs of the respective genes or gene fragments were cloned between *P_MET17_* and the coding sequence for CRU on the vector CRU pRS313 using E*ag*I and S*al*I restriction sites. N_ub_ fusions expressed from a centromeric plasmid were obtained by cloning the desired sequence in frame behind the N_ub_ coding sequence of the plasmid pCup1-Nui-HA kanMX. To fuse *mCHERRY* in front of *SEC4*, the ORF of *SEC4* was cloned behind the sequence of *P_CUP1_-mCHERRY* located on pCup-mCHERRY pRS313 using S*al*I and B*amH*I restriction sites. GFP or mCherry fusions to *SEC1, SNC1, and SSO1* were created by inserting the ORF of the respective genes behind the *P_MET17_-GFP/mCherry* module on a pRS315 vector using E*ag*I and S*al*I restriction sites. *GST* fusions were obtained by placing the ORF of the respective gene or gene fragment in frame behind the *E. coli GST* sequence on the pGEX-2T plasmid (GE Healthcare, Freiburg, Germany) using B*amH*I and E*coR*I restriction sites. Fusions to the human O6-Alkyl-DNA transferase (SNAP-tag, New England Biolabs, Beverly, MA) were expressed from plasmid pAGT-Xpress, a pET-15b derivative (Schneider et al., 2013). Gene fragments were inserted in frame into a multi-cloning site located between the upstream *6xHIS*-tag-coding sequence and the downstream SNAP-tag-coding sequence. The *6xHIS-tag* fusions were obtained by placing the ORF of the respective gene or gene fragment behind the *E. coli 6xHIS-tag* sequence on the pAC plasmid (Schneider et al., 2013). Lists of plasmids and yeast strains used in this study can be found in Tables S2 and S3 of the supplementary information. Plasmid maps can be obtained upon request.

### Growth conditions, cultivation of yeast strains and genetic methods

Genetic manipulations of yeast and cultivation in different media followed standard protocols (Guthrie and Fink 1991). Media for Split-Ub interaction tests contained 1 mg/ml 5-fluoro-orotic acid (5-FOA, Thermo Fisher Scientific, Waltham, MA, USA). All yeast strains used in this study were derivatives of JD47, a segregant from a cross of the strains YPH500 and BBY45 (Dohmen et al., 1995). Gene deletions were performed by one step PCR-based homologous recombinations using pFA6a-hphNT1, pFA6a-natNT2, pFA6a-kanMX6 and pFA6a-CmLEU2 as templates (Bähler et al., 1998; Janke et al., 2004).

### Split-Ub interaction analysis

Large-scale split-Ub assays were performed using a library of 389 different N_ub_ fusion-expressing yeast alpha-strains that were mated with a single a-strain expressing the CRU fusion. Matings, and replica plating on different media were performed with a RoToR HDA stamping robot (Singer Instruments, Somerset, UK) as described (Hruby et al., 2011; Neller et al., 2015). To individually measure interactions between single pairs of CRU and N_ub_ fusion proteins, JD47 cells expressing the respective CRU fusion protein were mated with JD53 cells expressing the respective N_ub_ fusion protein. The resulting diploid cells co-expressing both fusion proteins were spotted onto media containing or lacking 1 mg/ml FOA in 10-fold serial dilutions from OD_600_ = 1 to 0,001. The plates were grown at 30 °C and recorded every day for 4 to 5 days (Eckert and Johnsson, 2003).

### Preparation of yeast cell extracts

Exponentially grown yeast cell cultures were pelleted and resuspended in yeast extraction buffer (50 mM HEPES, 150 mM NaCl, 1 mM EDTA) with 1x protease inhibitor cocktail (Roche Diagnostics, Penzberg, Germany). Cells were lysed by vortexing them together with glass beads (3-fold amount of glass beads and extraction buffer to pellet weight) 12 times for one minute interrupted by short incubations on ice. The obtained yeast cell extracts were clarified by centrifugation at 16,000 g for 20 min at 4°C. Extracts were probed with anti-Pho8p (D3A10, Thermo Fisher Scientific, Waltham, MA, USA) or anti MYC antibodies, followed by goat anti mouse IGG (Sigma-Aldrich, Steinheim, Germany).

### Recombinant protein expression and purification from *E. coli*

All proteins were expressed in *E.coli* cells (BL21, Amersham, Freiburg, Germany). GST and all PH domains fused to GST were expressed at 30 °C for 5 h in LB medium, 1 mM IPTG. GST fusions to SH3 domains were expressed at 18 °C in SB medium with 0,2 mM IPTG over night. 6xHis-Sec1p was expressed over night at 18 °C in SB medium, 0,2 mM IPTG. 6xHis-Cdc42_G12V_ was expressed for 5 h at 30 °C in LB medium, 1 mM IPTG. 6xHis-Exo84_121-141_-AGT, 6xHis-Sec3_481-521_-AGT, and 6xHis-AGT were expressed at 30 °C in LB medium, 1 mM IPTG. Cells were pelleted, washed once with 1xPBS, resuspended in 1xPBS containing protease inhibitor cocktail (Roche Diagnostics, Penzberg, Germany) and lysed by lysozyme treatment (1 mg/ml, 30 min on ice), followed by sonication with a Bandelin Sonapuls HD 2070 (Reichmann Industrieservice, Hagen, Germany). Extracts were clarified by centrifugation at 40,000 g for 10 min at 4 °C. 6xHis-Sec1p and 6xHis-Cdc42_G12V_ were purified by metal affinity-(HisTrap, GE healthcare, Freiburg, Germany) followed by size exclusion chromatography (Superdex200, GE healthcare) in PBS or TRIS buffer (50 mM TRIS, 50 mM NaCl, 10 mM MgCl_2_, pH 8). 6xHis-Exo84_121-141_-AGT was purified by metal affinity-followed by anion exchange chromatography (ResourceQ, GE healthcare, Freiburg, Germany) in TRIS buffer (50 mM TRIS, 50 mM NaCl, pH 7,5). 6xHis-Sec3_481-521_-AGT was directly used after metal affinity chromatography and buffer exchange to PBS through PD10 column chromatography (GE Healthcare, Freiburg, Germany). AGT was purified by metal affinity chromatography and preparative size exclusion chromatography. Before use the monomeric and aggregate-free state of all purified 6x-His-tagged fusion proteins were verified through analytical size exclusion chromatography (Superdex600, GE Healthcare, Freiburg, Germany) in PBS.

### GST-pulldown assay

GST-tagged proteins were immobilized on glutathione sepharose beads (GE Healthcare, Freiburg, Germany) directly from *E. coli* extracts. After 1 h incubation at 4 °C with either yeast extracts or purified proteins under rotation at 4 °C, the beads were washed 3 times with the respective buffer. Bound material was eluted with GST elution buffer (50mM TRIS, 20mM reduced glutathione) and analyzed by SDS-PAGE followed by Coomassie staining and immunoblotting with anti-HIS (Sigma-Aldrich, Steinheim, Germany) or anti-MYC antibodies (Hruby et al., 2011).

### Liposome preparation and spin down assay

1,2-Dipalmitoyl-xn-glycero-3-phosphoserine (DPPS; Echelon Biosciences Inc., Salt Lake City, UT, USA) was dissolved in chloroform at a concentration of 10 mg/ml. For liposome preparation, lipids were dried down under a stream of nitrogen gas. From the resulting lipid film, a suspension of vesicles was generated using bath-type sonication in PBS. For spin down experiments lipids (final concentration: 0,25 mg/ml) and purified GST fusion proteins (final concentration: 0,25 μM) were mixed in a centrifuge tube and centrifuged for 1 h at 500 000 g at 4 °C. Supernatants and pellets were boiled in Lämmli-buffer and analyzed by western blotting with anti-GST antibodies (Sigma-Aldrich, Steinheim, Germany).

### Transmission electron microscopy

*Δboi2 P_GAL1_BOI1*-cells (with or without additional plasmids) were grown for 10 or 12 h in liquid SD media containing either galactose or glucose. Cells were washed 3 times with PBS and fixed with 2 % of glutaraldehyde (1h) followed by 2 % osmiumtetroxyde (chemPur GmbH, Karlsruhe, Germany). After dehydration in increasing ethanol concentrations (30%, 50%, 70%, and 90%) cells were incubated with uranyl acetate (Merck, Darmstadt, Germany) for 30 min.. After 3 wash steps with ethanol, samples were incubated in increasing concentrations of Spurr resin solved in propanol (2:1, 1:2, 1:3 for 1h each step) (Polysciences Inc., Warminster, PA, UA). Afterwards, the samples were incubated in Spurr resin over night. Resin polymerization occurred at 60 °C for 48 hours. Ultrathin sections (about 70 nm) were cut with a Leica Ultracut UCT ultramicrotome using a diamond knife (Diatome, Biel, Switzerland). Sections were analyzed with a Jeol 1400 TEM (Jeol, Tokyo, Japan) and the images were digitally recorded with a Veleta camera (Olympus, Münster, Germany).

### Fluorescence microscopy

Fluorescence microscopic images were generated with an Axio Observer spinning disc confocal microscope (Zeiss, Göttingen, Germany) equipped with an Evolve512 EMCCD camera (Photometrics, Tucson, USA), a Plan-Apochromat 63X/1.4 Oil DIC objective, and 488 nm and 561 nm diode lasers (Zeiss, Göttingen, Germany). Operations were performed with the ZEN2012 software package (Zeiss). SPLIFF analysis was performed with a DeltaVision system (GE Healthcare, Freiburg, Germany) provided with an Olympus IX71 microscope (Olympus, Münster, Germany) equipped with a CoolSNAP HQ^2^-ICX285 or a Cascade II 512 EMCCD camera (Photometrics, Tucson, USA), a 100×UPlanSApo 100×1.4 Oil ∞/0.17/FN26.5 objective (Olympus, Münster, Germany), a steady state heating chamber, and a Photofluor LM-75 halogen lamp (Burlington, VT, USA).

Yeast cells were prepared for microscopy as previously described (Schneider et al., 2013). In brief, an overnight culture (grown in liquid SD medium) was diluted 1:15 in 3 ml fresh SD medium and incubated for 3 to 5 h to mid-log phase. Cells were spun down and resuspended in 50 μl fresh medium. 3 μl of this suspension were transferred to a microscope slide, covered with a glass coverslip and analyzed under the microscope. For time-lapse analysis, cells were immobilized with a coverslip on custom-designed glass slides containing solid SD medium with 1,8 % agarose.

### Quantitative analysis of microscopy pictures

Microscopy files were analyzed and processed using ImageJ64 1.45s (US National Institute of Health). All images were acquired as adapted z-series and projected to one layer. For the quantitative comparison of the mean fluorescent intensities in daughter and mother cells (I_D_/I_M_), randomly selected areas around the cells were measured as background and subtracted from the respective intracellular intensities. mCherry-Sec4p fluorescence intensity profiles were measured using the ImageJ plot profile tool. All fluorescence quantifications were performed with z-overlays of seven stacks.

### SPLIFF interaction analysis

Analysis of temporal and spatial characteristics of the Boi1p/Sec1p interaction by SPLIFF was performed as described (Moreno et al., 2013; Dünkler et al., 2015). For SPLIFF interaction measurements a-cells (*P_MET17_-BOI1CCG*) and alpha-cells (*N_ub_-SEC1, N_ub_-PTC1*) cells were grown in SD media without methionine and 5 μM Cu. Equal amounts of cells were mixed shortly before agarose slide preparation. Image acquisition was started as soon as the first mating projections appeared. Zygote formation was recorded by three-channel z-stacks (5 × 0.6 μm distance) every 2 min. 2D Images of the GFP- and mCherrry channel were created by SUM-projections for each time point. To measure the fluorescence signals over time the integrated density of the region of interest and a region within the cytosol was determined. The cytosolic value was subtracted from the localized signal to obtain the localized fluorescence intensity for each channel (FI_red_ and FI_green_) and normalized to the values shortly before cell fusion. The resulting relative fluorescence intensity RFI(t) was applied to calculate the conversion FD(t):

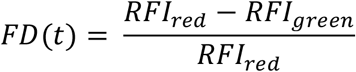

### Latrunculin A treatment and actin phalloidin staining

200 μl exponentially growing yeast cells were treated for 12 min under shaking with 2 μl of a 10 mM latrunculin A (Sigma-Aldrich GmbH, Steinheim, Germany) stock solution dissolved in DMSO. Afterwards, cells were fixed with a final concentration of 3,7 % formaldehyde. Cells were washed in PBS and incubated with Alexa-Fluor 594 phalloidin (Thermo Fisher Scientific, Waltham, MA, USA) at a final concentration of 66 nM for 30 min at 4 °C.

### Tetrad analysis

Tetrad analysis was performed as described (Neller et al., 2014). To prove the synthetic lethality between *BOI1* and *BOI2*, deletions of each gene were generated in the haploid strains JD47 and JD53. Afterwards Δ*boi1*- and Δ*boi2*-cells were mated to generate heterozygous diploids. After sporulation, asci were dissected and single spores spotted onto YPD agar plates (MSM300; Singer Instruments, Somerset, UK) and grown for 4-6 days at 30 °C. The allelic compositions of the spores were analyzed by exposing the cells to suitable selection media and by performing diagnostic PCRs.

### SPR measurement

To measure the K_D_ value of the 6xHIS-Exo84_121-141_-AGT/6xHIS-SH3_Boi1_ complex surface plasmon resonance measurements using a Biacore X100 system (GE Healthcare, Freiburg, Germany) were performed as described (Renz et al., 2013). In brief, purified 6xHIS-Exo84_121-141_-AGT, covalently labeled with SNAP-Biotin (New England Biolabs) in HBSEP buffer (100 mM HEPES, 1.5 M NaCl, 30 mM EDTA, 0.5% (w/v) Tween20; pH 7.4) was captured on the surface of a CM5 chip (GE Healthcare, Freiburg, Germany) previously coated with an anti-Biotin antibody (US Biologicals, Pittsburgh, PA, USA). The chip was titrated with increasing concentrations of 6xHIS-SH3_Boi1_ in HBSEP buffer. The data were analyzed with the Biacore Evaluation Software (Version 1.1; GE Healthcare, Freiburg, Germany). The sensor chip was regenerated with 12 mM NaOH after each experiment.

### Statistical evaluation

Statistical data evaluations were performed using GraphPad Prism5. T-tests were used to compare the percentage of cells of a certain phenotype. Mann-Whitney-U-tests were applied to evaluate the significance of the differences between ratios of the mean fluorescent intensities in daughter and mother cells of different genotypes.

## Acknowledgements

We thank Steffi Timmermanns, and Ute Nussbaumer for technical assistance. We thank Reinhardt Fischer (KIT, Karlsruhe, Germany) for *A. nidulans* genomic DNA.

## Competing interests

The authors declare no competing financial interests.

## Funding

The work was funded by grants from the Deutsche Forschungsgemeinschaft (DFG) (Jo 187/5-2; Jo 187/8-1). J.K was supported by a fellowship from the Graduate School in Molecular Medicine, Ulm University.

